# Plum is a novel regulator of synaptic function and muscle size in *D. melanogaster*

**DOI:** 10.1101/463885

**Authors:** Hrvoje Augustin, Jereme G. Spiers, Nathaniel S. Woodling, Joern R. Steinert, Linda Partridge

## Abstract

Alterations in the neuromuscular system underlie several neuromuscular diseases and play critical roles in the development of sarcopenia, the age-related loss of muscle mass and function. Mammalian Myostatin (MST) and GDF11, members of the TGF-β superfamily of growth factors, are powerful regulators of muscle size in both model organisms and humans. Myoglianin (MYO), the *Drosophila* homolog of MST and GDF11, is a strong inhibitor of synaptic function and structure at the neuromuscular junction (NMJ), and a negative regulator of body weight and muscle size and function in flies. Here, we identified Plum, a cell surface immunoglobulin homologous to mammalian developmental regulators Protogenin and Nope, as a modulator of MYO function in the larval neuromuscular system. Reduction of Plum specifically in the larval body-wall muscles abolishes the previously demonstrated positive effect of attenuated MYO signalling on both muscle size and neuromuscular junction structure and function, likely by de-sequestrating the remaining MYO. In addition, downregulation of Plum on its own results in decreased synaptic strength and body weight, classifying Plum as a (novel) regulator of neuromuscular function and body (muscle) size. These findings offer new insights into possible regulatory mechanisms behind ageing- and disease-related neuromuscular dysfunctions in humans and identify potential targets for therapeutic interventions.

## INTRODUCTION

The neuromuscular system is comprised of individual motor units, each consisting of a single motor neuron, a neuromuscular junction and all the muscle fibres innervated by the motor neuron. Diminished motor unit function and decreased muscle volume are hallmarks of several neuromuscular disorders (Gonzalez-Freire et al., 2014) and of sarcopenia, the age-associated loss of skeletal muscle mass and function (Cruz-Jentoft et al., 2010). Gradual loss of skeletal muscle capacity has been reported in invertebrates, rodents, and humans (Eddinger et al., 1985; Herndon et al., 2002; Young et al., 1985), with the intrinsic mechanisms regulating age-related muscle dysfunction largely conserved across species (Demontis et al., 2013). Age-related muscle loss is accompanied by progressive modifications in the structure and function of the neuromuscular junction (NMJ), the specialised synapse at the interface between the nervous and muscular system (Gonzalez-Freire et al., 2014), with the resulting uncoupling of the excitation–contraction machinery (Delbono, 2000; Delbono et al., 1995). In mammals, including humans, these modifications include changes in the branching pattern of the motor nerve terminal that contacts the myofiber, fragmented NMJ architecture, impaired synaptic acetylcholine receptor distribution, and decreased density of presynaptic active zone markers (Barker and Ip, 1966; Chen et al., 2012; Nishimune et al., 2016; Oda, 1984; Wokke et al., 1990). Functionally, aged mammalian NMJs exhibit increased failures in evoked release (Lee et al., 2017), changes in quantal release (Kelly and Robbins, 1983) and a slowing-down of axoplasmic transport of proteins (Gutmann and Hanzlikova, 1973). Skeletal muscle and NMJ deficits are found in many motoneuronal and neuromuscular disorders, with impaired neurotransmission and muscle wasting characterising amyotrophic lateral sclerosis (ALS) (Cappello and Francolini, 2017; Chand et al., 2018; Pansarasa et al., 2014), myotonic and muscular dystrophies (Bombelli et al., 2016; Malatesta, 2012; Mateos-Aierdi et al., 2015; Rudolf et al., 2014; Shin et al., 2013), myasthenia gravis (Fambrough et al., 1973; Shigemoto et al., 2010) and others. Whether muscle loss precedes or follows the changes in the function of the NMJ is currently unresolved, but animals studies suggest that NMJ remodelling might play a critical role in the progression of sarcopenia (Ezaki et al., 2000).

*Drosophila melanogaster* is a convenient and proven model system for studying various aspects of developmental regulation of muscle mass and control of NMJ function (Augustin and Partridge, 2009; Kreipke et al., 2017; Plantie et al., 2015). *Drosophila* larval glutamatergic NMJs share structural and functional similarities with mammalian cholinergic junctions (Budnik, 1996) and striated muscles in *Drosophila* resemble vertebrate skeletal muscles in structure, function, and protein composition (Augustin and Partridge, 2009). Previously, we used this model system to investigate the role of *Drosophila* MYO, the muscle- and glia-expressed fly homolog of TGF-β growth factors Myostatin (MST) and GDF11 (Lo and Frasch, 1999), in regulating synaptic function, muscle size and body weight (Augustin et al., 2017). MST (also known as GDF-8) is a circulating cytokine that serves as a powerful negative regulator of muscle mass in mammalian species (Carnac et al., 2007; McPherron et al., 1997). In addition to its MST-like role as an inhibitor of larval weight and muscle size, muscle-derived MYO is a strong negative regulator of neurotransmission, synaptic morphology and the density of critical pre- and post-synaptic components (Augustin et al., 2017).

Plum is a *Drosophila* transmembrane, immunoglobulin superfamily protein (Yu et al., 2013) and a distant homolog of Protogenin and Nope, mammalian regulators of developmental processes in nervous, muscle and epithelial tissues (Salbaum and Kappen, 2000; Takahashi et al., 2010; Wong et al., 2010). Protogenin was also associated with attention deficit hyperactivity disorder (Wigg et al., 2008) and Nope is a surface marker for human and murine liver cancers (Marquardt et al., 2011; Xiang et al., 2016). Recently, Plum was identified as a modulator of axon pruning in the *Drosophila* nervous system (Yu et al., 2013). Plum genetically interacts and interferes with MYO function, likely by physically sequestrating MYO (Yu et al., 2013). In this study, we examined the interactions between Plum and MYO in regulating larval muscles and NMJ physiology. We identified Plum as a novel modulator of MYO action on NMJ synaptic transmission and muscle size, and an independent regulator of synaptic strength and larval weight.

## RESULTS

### Muscle-derived Plum regulates NMJ synapse strength independently and by modulating MYO

We previously showed that genetic attenuation of *Myo* specifically in the larval somatic muscle, using a microRNA construct to target the *Myo* transcript (genotype: *Mef2-GAL4/UAS-miRNAmyo*) (Awasaki et al., 2011), increases muscle size, NMJ synaptic transmission and locomotion by >20% (Augustin et al., 2017), defining MYO as a potent neuromuscular inhibitor in flies. Considering the large size of the (postsynaptic) muscle compartment relative to the (presynaptic) motoneuronal and glial compartments, and the expression of the Plum mammalian homolog Nope in developing skeletal muscles (Salbaum and Kappen, 2000), we investigated interactions between Plum and MYO by knocking-down Plum in the larval body-wall muscle. We therefore analysed phenotypes in double knock-down MYO-Plum larvae (*Mef2-GAL4/UAS-miRNAmyo/plumRNAi*) and single knock-down Plum animals (*Mef2-GAL4/UAS-plumRNAi*).

We first examined the impact of Plum down-regulation on NMJ physiology. Body-wall muscles in developing flies consist of bilaterally symmetrical hemi-segments composed of 30 multinucleated muscle fibres (Ruiz-Canada and Budnik, 2006). We focused our analyses on muscles 6 and 7, large myofibres innervated by two motoneurons forming a single, excitatory, chemical (glutamatergic) NMJ. Contractions of these muscles are triggered by “non-spiking” postsynaptic potentials that are graded in duration and amplitude, allowing for quantitative comparisons between genotypes (Peron et al., 2009). The amplitude of these Ca^2+^-dependent, nerve-evoked postsynaptic excitatory junctional currents (eEJCs) reflects either the magnitude of presynaptic release or the postsynaptic sensitivity to neurotransmitter (Peron et al., 2009). Muscle-specific reduction of MYO leads to dramatically increased evoked response (Augustin et al., 2017); simultaneous suppression of *Plum*, however, reversed the response to control (*+Mef2-GAL4*) levels, with the down-regulation of *Plum* alone further diminishing the evoked currents. Further examination of the data showed a statistically significant interaction between the effects of simultaneously reduced MYO and Plum relative to the control genotype (two-way ANOVA analysis in *Fig. 1B*). We then measured the amplitudes of spontaneous “miniature” postsynaptic currents (mEJCs), also known as “quantal size” (Petersen et al., 1997). The mEJCs represent postsynaptic responses to a single presynaptically released vesicle containing neurotransmitter and are a reliable indicator of the density of functional, NMJ, glutamate receptors (DiAntonio et al., 1999). While the mean mEJC amplitude did not differ significantly between genotypes (*Figs. 1A* and *1C*), the frequency distribution analysis revealed that Plum down-regulation in either control or reduced MYO background caused a strong shift toward smaller “miniature” currents (*Fig. 1D*). The number of presynaptic release sites or probability of spontaneous vesicle fusion events was not affected, as demonstrated by the normal mEJC frequency in mutant genotypes (*Fig. 1E*). Taken together, our electrophysiological results imply that MYO and Plum affect NMJ physiology by controlling the density of the postsynaptic glutamate receptor field, with Plum having a modulatory effect on MYO and acting as an autonomous synaptic regulator.

**Figure 1.**
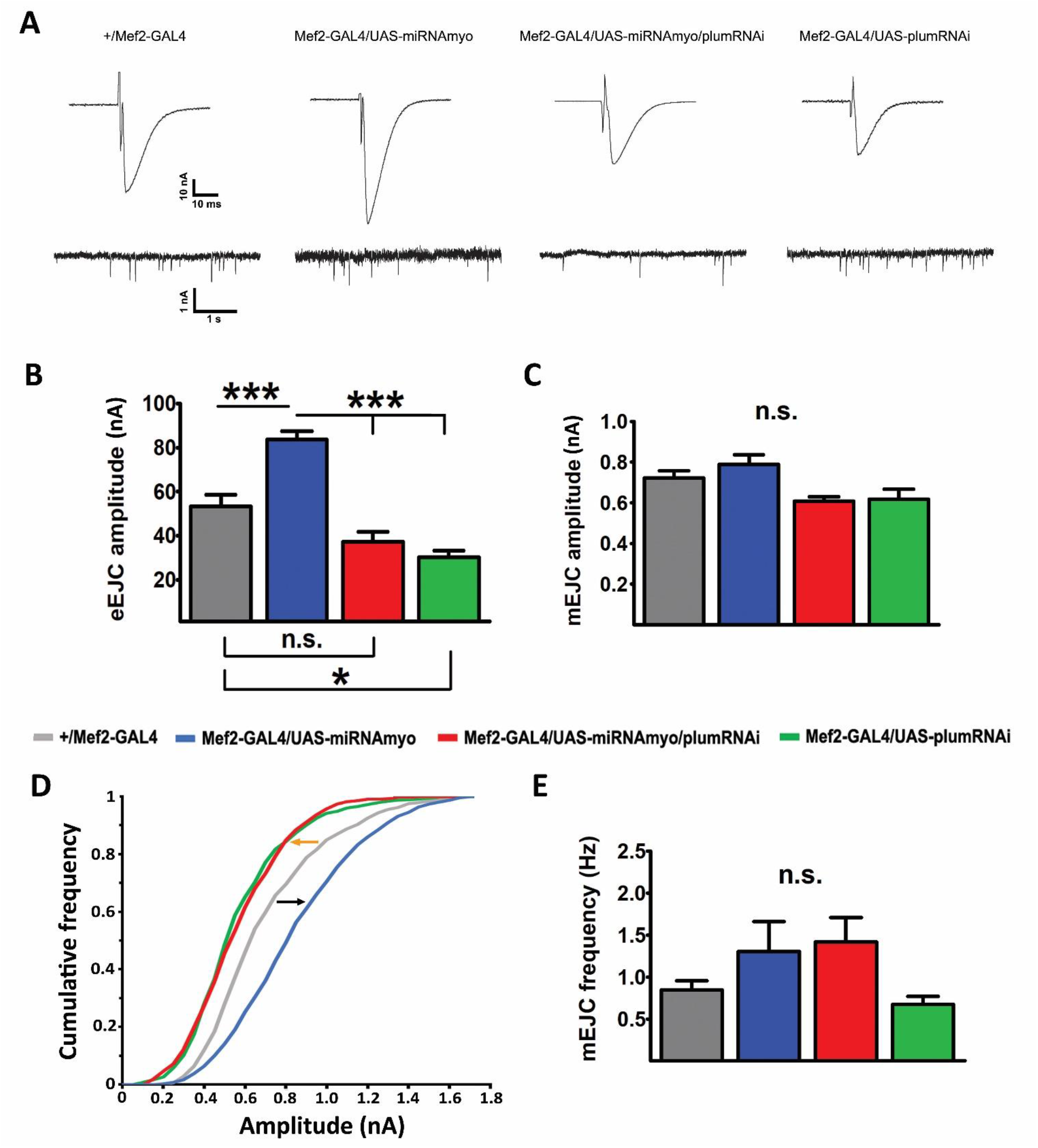
Plum is a regulator of neurotransmission at the larval NMJ. (*A*) Representative traces of evoked (top) and spontaneous (“miniature”) synaptic responses recorded from muscle 6. Histograms of evoked (*B*) and spontaneous (*C*) responses for given genotypes (control: *+/Mef2-GAL4*; MYO downregulation: *Mef2-GAL4/UAS-miRNAmyo*; MYO and Plum down-regulation: *Mef2-GAL4/UAS-miRNAmyo/plumRNAi*; Plum down-regulation: *Mef2-GAL4/UAS-plumRNAi*) (*n* = 7-16). Two-way ANOVA analysis: the interaction between *Mef2-GAL4/UAS-miRNAmyo and Mef2-GAL4/UAS-miRNAmyo/plumRNAi* is highly significant (P<0.0001). (*D*) Cumulative frequency distribution diagram of mEJC amplitudes. Black arrow indicates shift toward higher amplitude “minis”, orange toward lower. For all paired analyses except between *Mef2-GAL4/UAS-miRNAmyo/plumRNAi* and *Mef2-GAL4/UAS-plumRNAi* KS test, P<0.0001 (n=7-18 animals, ~1500-2000 events measured per genotype). (*E*) Histogram of mEJC frequencies (*n* = 7-13). Bar graphs: error bars indicate SEM (one-way ANOVA + Tukey’s post-test: *p < 0.05, ***p < 0.001, n.s. = not significant).

### Plum modulates the action of MYO on synapse structure and receptor composition

Postsynaptic receptors at the larval NMJ are AMPA-type tetrameric complexes formed by glutamate receptor (GluR) subunits IIC, IID and IIE, in addition to either subunit IIA or IIB (Featherstone et al., 2005; Qin et al., 2005). Assemblies containing the IIA subunit (GluRIIA) are pharmacologically and biophysically distinct from the ones incorporating GluRIIB and carry the bulk of the ionic current at this synapse (DiAntonio et al., 1999; Petersen et al., 1997). We have recently shown that elevated evoked and spontaneous synaptic currents in “low MYO” larvae show corresponding increase in the density of GluRIIA receptors (Augustin et al., 2017), in line with previously demonstrated correlation between GluRIIA levels and either evoked response (Sigrist et al., 2002) or quantal size (DiAntonio et al., 1999). We therefore used immunohistochemistry to measure the area of GluRIIA clusters in the NMJ boutons (*Fig. 2A*) of control and mutant animals. The GluRIIA cluster area is directly proportional to the number of functional GluRs measured electrophysiologically, and independent of changes in NMJ morphology (Featherstone et al., 2002; Liebl and Featherstone, 2008; Rasse et al., 2005).

**Figure 2.**
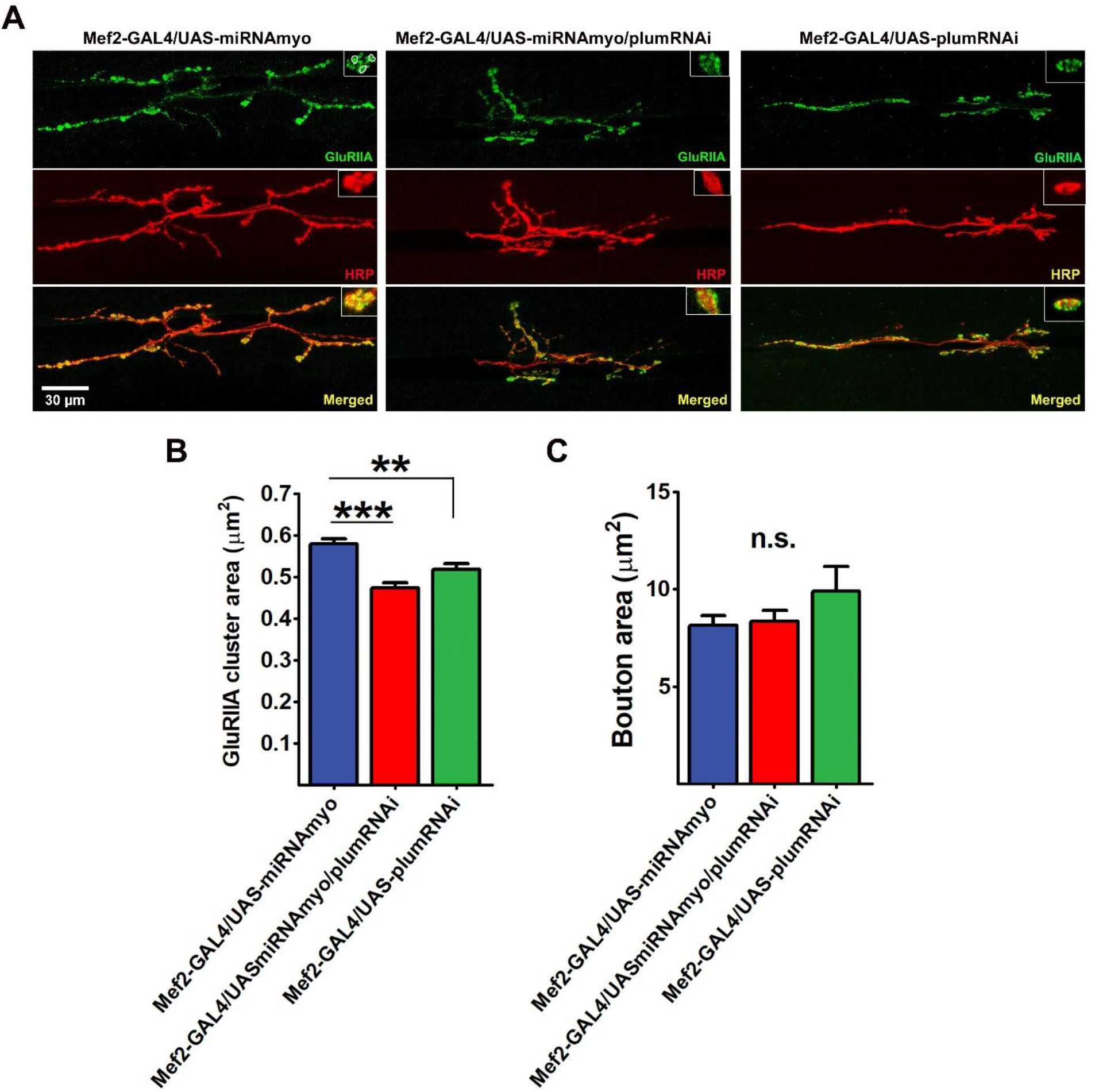
Downregulation of *plum* reduces the size of GluRIIA clusters in reduced MYO background. (*A*) Representative confocal images of the 3rd instar larval 6/7 NMJs in denoted genotypes. Insets show synaptic varicosities (boutons), with individual GluRIIA clusters within boutons marked by circled areas. Anti-HRP antibody visualised the presynaptic (motoneuronal) membrane. (*B*) Quantification of GluRIIA cluster areas (n = 8-9). (*C*) The mean size of synaptic boutons was not affected by *plum* downregulation (n = 7-9). All bar graphs: error bars indicate SEM (one-way ANOVA + Tukey’s post-test: **p < 0.01, ***p < 0.001, n.s. = not significant).

Our results showed that *Plum* down-regulation in the muscle led to significantly (~20%) smaller GluRIIA clusters in animals with muscle-reduced *Myo* expression (*Figs. 2A-B*). This reduction was identical in magnitude to the increase in the density of IIA-type receptors upon MYO downregulation relative to the control genotype (Augustin et al., 2017), demonstrating the reversal to wild-type receptor levels caused by reduced Plum. Unlike its effect on evoked response and distribution of mEJC amplitudes, muscle-specific *Plum* suppression alone was unable to further reduce the GluRIIA cluster area. The lack of perfect correlation between electrophysiological analyses and antibody staining likely occurred because the latter cannot distinguish between functional and non-functional glutamate receptors. Down-regulation of Plum and/or MYO in the muscle had no effect on the size of NMJ boutons (*Fig. 2C*), suggesting that the density of GluR clusters did not depend on spatial constraints within boutons.

Our immuno-staining data confirm the negative effect of Plum down-regulation on enhanced neurotransmission caused by muscle-specific knock-down of MYO.

### The number of Brp puncta scales with NMJ size upon Plum and/or MYO attenuation

Bruchpilot (Brp) is a presynaptic marker at the larval NMJ and the *Drosophila* homolog of the vertebrate active zone protein ELKS (Wagh et al., 2006). Brp is required for function and structural integrity of synaptic active zones and is necessary for normal evoked, but not spontaneous, release at the glutamatergic synapse of the NMJ (Wagh et al., 2006).

We have previously shown that enhanced evoked response in reduced MYO background correlates with the increase in the NMJ size (Augustin et al., 2017). These findings agree with the previously established positive correlation between the number of boutons and the strength of evoked signal transmission (Sigrist et al., 2003; Sigrist et al., 2002). While the number of active zones (measured as the number of distinct Brp puncta) per bouton is unchanged between genotypes (*Figs. 3A-B*), the reversal of the increased amplitude of evoked synaptic responses in reduced MYO larvae upon Plum suppression (*Fig. 1B*) could be explained by the reduced bouton number, NMJ branching, and NMJ length (*Fig. 3C*) in the larvae suppressing both MYO and Plum. In addition, Plum likely exhibits a broader, non-Brp related, physiological effect because the morphological changes alone cannot explain the negative effect of *Plum* knock-down on synaptic strength in the control background (green bar in *Fig. 1B*). These findings implicate Plum as a modulator of synaptic strength in low MYO background via its impact on NMJ morphology.

**Figure 3.**
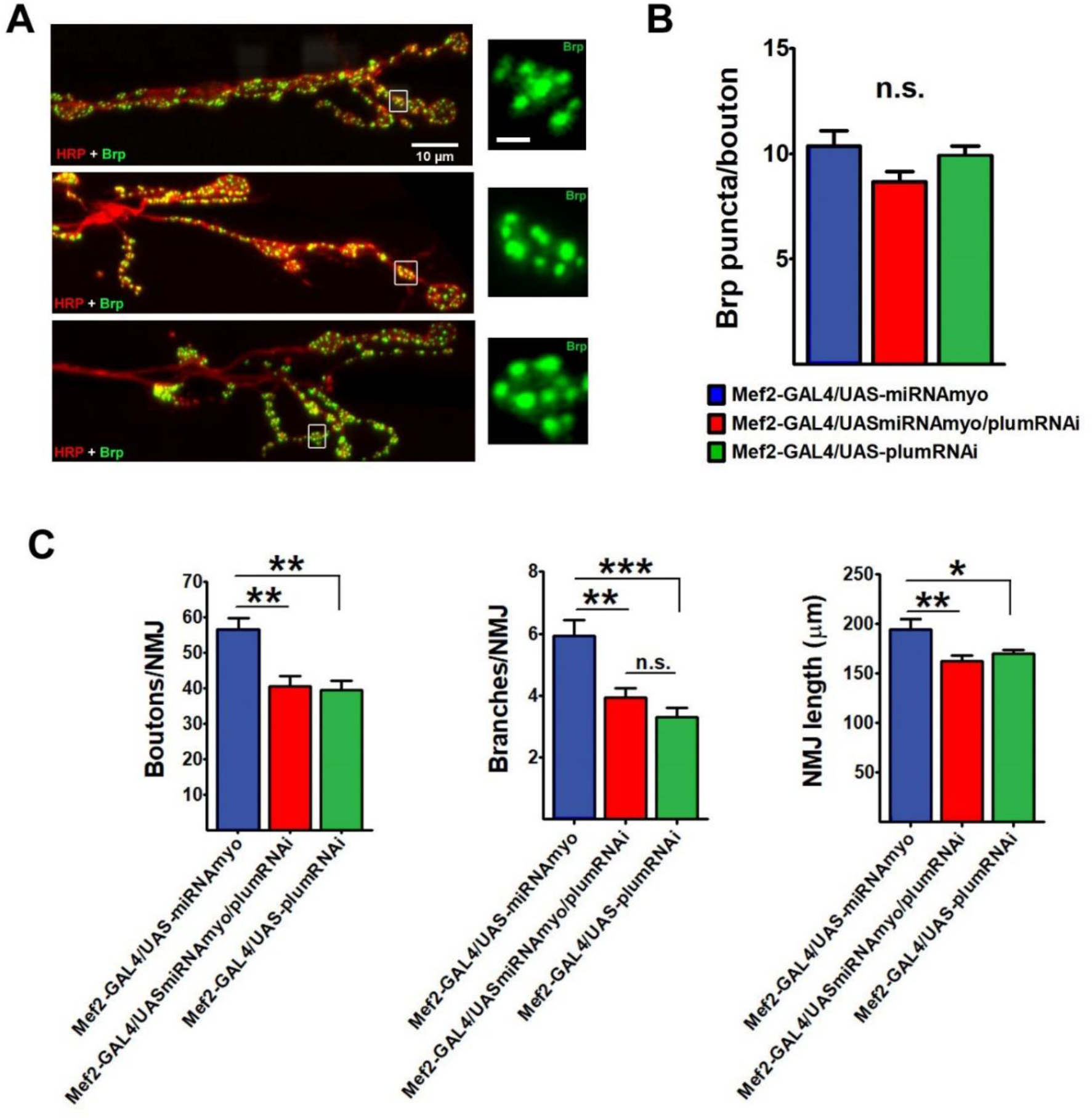
Number of Brp puncta and NMJ morphology. (*A*) Representative confocal images of distal NMJ segments. Insets show Brp puncta in individual boutons (inset scale bar: 1 μm). (*B*) Mean number of Brp puncta per bouton (n = 7-9). (*C*) NMJ morphology: the number of boutons (left) and branches (middle) per 6/7 NMJ and NMJ length (right) (n = 9-17). All bar graphs: error bars indicate SEM (one-way ANOVA + Tukey’s post-test: *p < 0.05, **p < 0.01, ***p < 0.001, n.s. = not significant).

### Knock-down of *Plum* abolishes the effect of reduced MYO on muscle size and body weight

Muscle-derived MYO negatively regulates larval weight and muscle size (Augustin et al., 2017). We examined the effect of *Plum* attenuation on the size of body-wall muscles 6 and 7 (*Fig. 4A*) and on total larval body weight in reduced MYO background. Plum suppression completely abolished the positive effect of MYO down-regulation on the combined area of muscles 6 and 7 (*Fig. 4B*) and on larval wet weight (*Fig. 4C*), mirroring its effects on synaptic physiology (*Fig. 1*), synaptic composition (*Fig. 2B*) and NMJ morphology (*Fig. 3C*). The interaction between *MYO* and *Plum* down-regulation was significant, indicating a combinatorial effect of these interventions in abolishing the positive effect of reduced MYO on muscle size and body weight (two-way ANOVA analysis in *Figs. 4C-D*). Furthermore, Plum has an independent effect on body mass, because *Plum* knock-down larvae exhibit significantly reduced wet weight (*Fig. 4C*, green bar).

**Figure 4.**
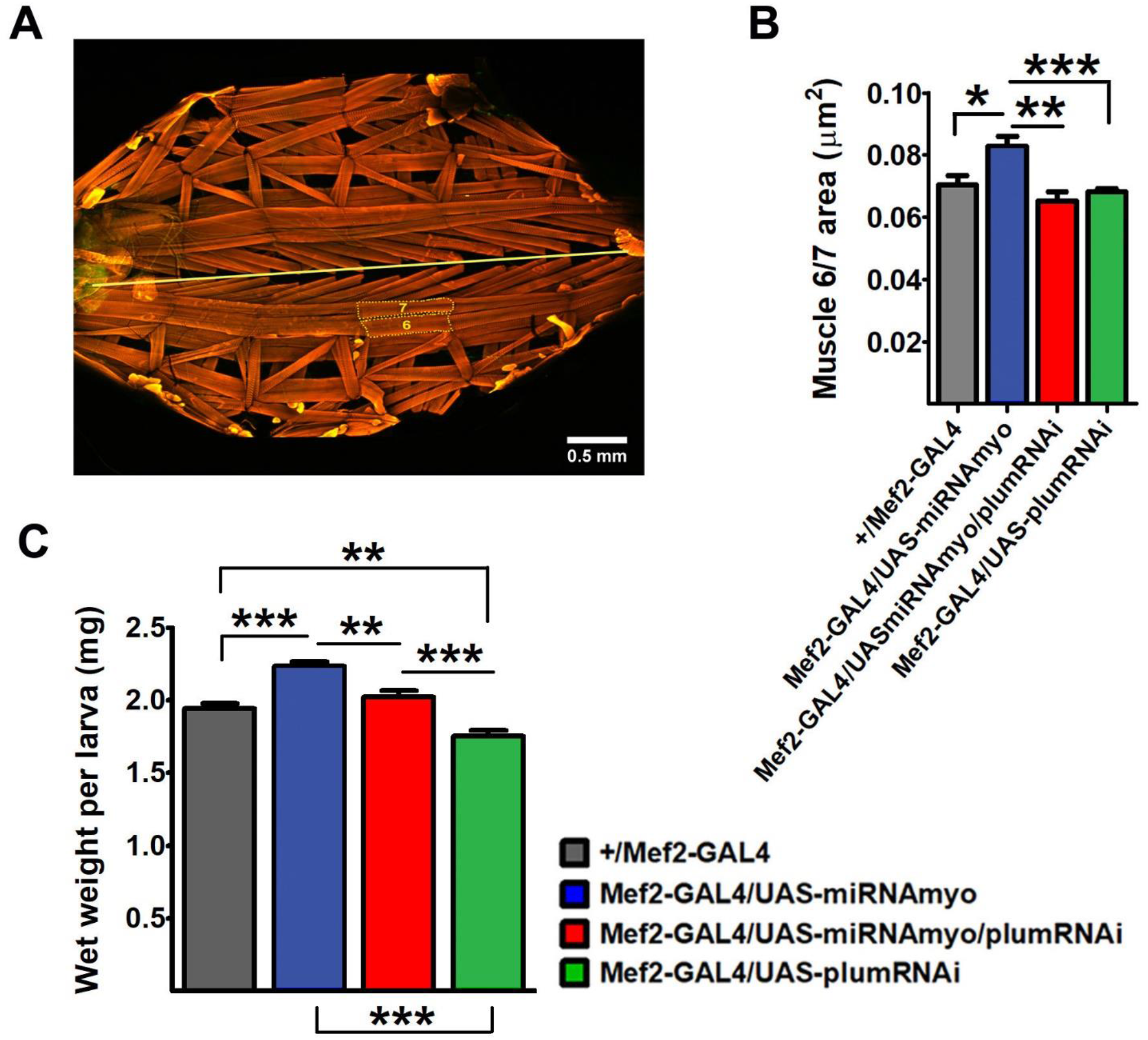
Plum regulates muscle size and larval weight. (*A*) Third instar larval preparation showing body wall muscles stained with phalloidin (anterior is on the left). Straight yellow line marks the midline. Dotted lines mark muscles 6 and 7. (*B*) Bar graphs compare mean combined area of muscles 6 and 7. (*C*) Total larval wet weight in indicated genotypes. In both (*B*) and (*C*), two-way ANOVA analysis indicates statistically significant interactions between *Mef2-GAL4/UAS-miRNAmyo and Mef2-GAL4/UAS-miRNAmyo/plumRNAi* (P=0.0191 and P=0.0040, respectively). Error bars in the bar graphs indicate SEM (one-way ANOVA + Tukey’s post-test: *p < 0.05, **p < 0.01, ***p < 0.001).

These experimental results identify Plum as a critical modulator of the action of MYO on the neuromuscular physiology, muscle size and weight, and as an independent regulator of synaptic strength and body weight in developing *D. melanogaster*.

## DISCUSSION

Chemical transmission across the neuromuscular junction is critical for converting action potentials originating in the central nervous system into contractile activity in skeletal muscles. It is therefore not surprising that impairment of the NMJ, in many cases, leads to reduced muscle mass and function in both vertebrates and invertebrates (Augustin and Partridge, 2009; Tintignac et al., 2015). In addition to motor unit elimination (Piasecki et al., 2016), reduced motor axon conduction velocity (Metter et al., 1998), diminished motor cortex excitability (Clark et al., 2014; Oliviero et al., 2006), and modified activity of muscle-intrinsic factors (Proctor et al., 1998), NMJ dysfunction is strongly correlated with decreased skeletal muscle size and strength under both healthy and pathological conditions (Pratt et al., 2015). For example, recent studies suggested that the malfunction of the NMJ plays a causative role in the onset of sarcopenia (Butikofer et al., 2011), and proposed NMJ stabilisation as a way to delay its progression (Mori et al., 2014). In dystrophic *mdx* mice, both pre- and post-synaptic abnormalities in the NMJ contribute to reduced muscle contractility (Pratt et al., 2015) and therapeutic approaches that specifically target NMJs have been proposed for treating spinal muscular atrophy and, possibly, ALS (Boido and Vercelli, 2016).

Several mammalian proteins have identified roles in linking structural and functional properties of the NMJ and (skeletal) muscles, the most important being Agrin and MuSK (neither of which has an obvious homolog in *D. melanogaster*). A cleaved fragment of the proteoglycan Agrin was identified as a marker for the diagnosis of sarcopenia (Hettwer et al., 2013) and implicated in the pathogenesis of sarcopenia caused by degeneration of the NMJ (Drey et al., 2013). Degradation of Agrin results in structural changes in the NMJ and innervated muscles, consistent with the notion that impaired NMJ functionality plays a role in the onset of sarcopenia (Butikofer et al., 2011). Agrin was first discovered as a neurotrophic factor sufficient for pre- and post-synaptic NMJ assembly and stabilisation (Bezakova and Ruegg, 2003) and aggregation of the junctional acetylcholine receptors (AChRs) (Ferns et al., 1993). The clustering of AChRs and postsynaptic differentiation is initiated by the release of Agrin from the innervating motor axon, which in turn activates muscle-expressed tyrosine kinase MuSK (Muscle-specific kinase) (DeChiara et al., 1996). Reduced function and density of synaptic AChRs at the NMJ is a hallmark of “normal” human (Oda, 1984) and rodent (Li et al., 2011) ageing muscles, and of several NMJ disorders characterised by skeletal muscle loss and weakness (Kong et al., 2009; Querol and Illa, 2013; Xu and Salpeter, 1997). These findings underline the importance of investigating links between the NMJ and muscle function and searching for novel regulators of these processes.

Just like Agrin, *Drosophila* MYO is a secreted ligand that regulates nervous system development (Awasaki et al., 2011) and NMJ synaptogenesis (Augustin et al., 2017). While MYO of glial origin governs remodelling of mushroom body neurons in developing animals (Awasaki et al., 2011), muscle-derived MYO functions as strong negative regulator of the size of larval somatic muscles and NMJ size and function (Augustin et al., 2017; Yu et al., 2013). In this study, we expanded our investigation of the role MYO in the larval neuromuscular system by analysing its interaction with Plum, a trans-membrane protein recently found to modulate MYO signalling during mushroom body development (Yu et al., 2013). Specifically, Plum sequestrates MYO, inhibiting its pruning-promoting effect on mushroom body neurons, with the reduction of endogenous Plum stimulating pruning (Yu et al., 2013). The “low MYO” animals used in our experiments have the *Myo* transcript levels reduced by ~60% (Augustin et al., 2017), with the remaining circulating MYO possibly sequestered by Plum. De-sequestering this Plum-bound MYO could therefore reverse some of the effects of genetic MYO attenuation on NMJ function and muscle size. In agreement with this hypothesis, simultaneous MYO and Plum suppression in the muscle completely reversed the effect of reduced MYO on synaptic physiology, muscle size, and total body weight. In addition, Plum reduction alone (i.e. without concomitant down-regulation of MYO) reduced the synaptic strength and body weight below control levels (*Fig. 5*), identifying muscle-derived Plum as a novel and independent target for manipulating neuromuscular function. It is important to note that MYO and Plum regulate physiological properties of the NMJ synapse by controlling the number of glutamate receptors, the functional analogues of ACh receptors in mammalian NMJs.

**Figure 5.**
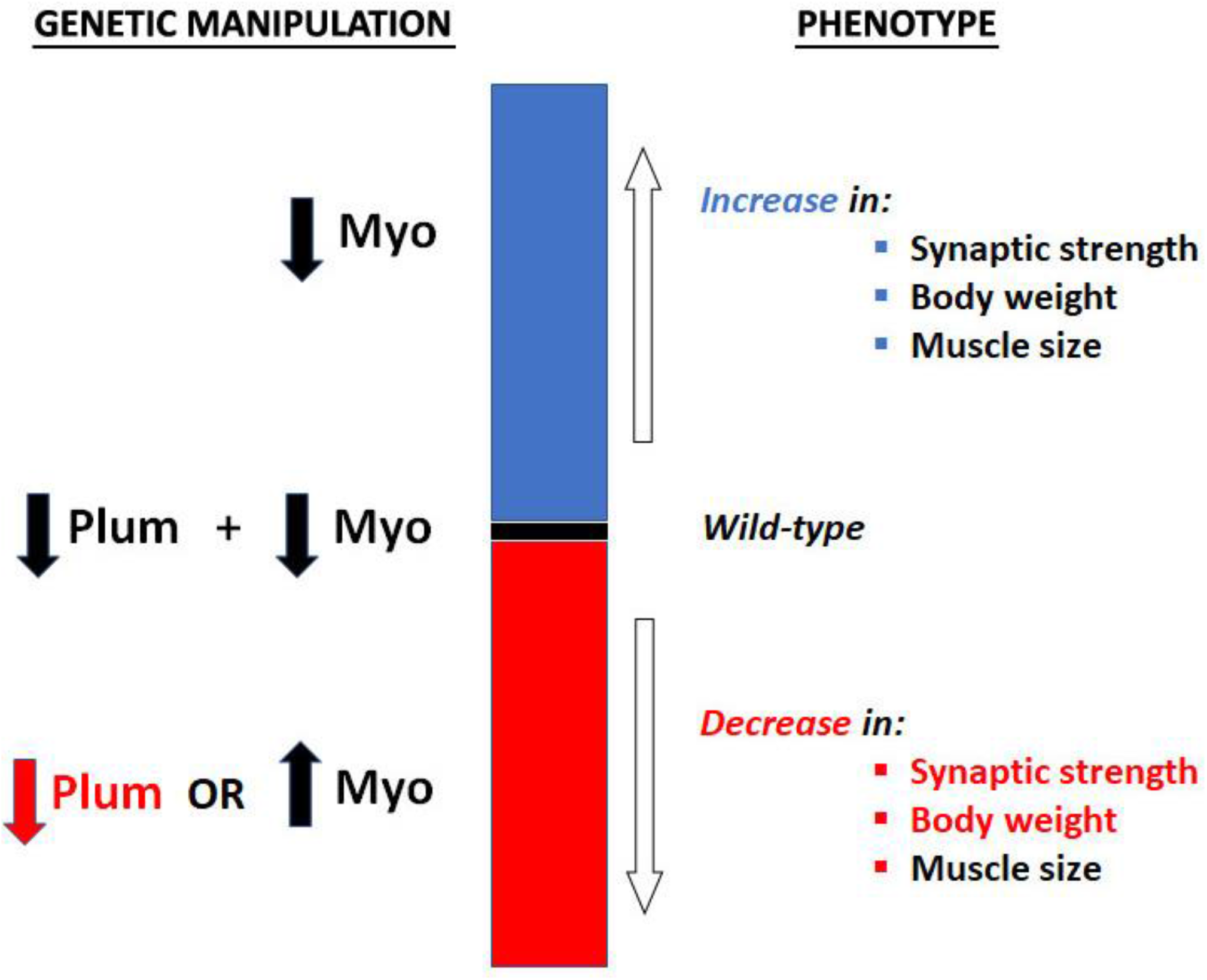
Diagram illustrating the effects of MYO and Plum on synaptic strength, muscle size and body weight in *Drosophila* larvae. Plum reduction abolishes the (positive) effect of reduced MYO on synaptic strength, total body weight and muscle size. Attenuation of Plum on its own, or MYO up-regulation, negatively affect synaptic strength and larval weight, with MYO over-expression having an additional (negative) effect on muscle size.

Sequestration is a well-described mechanism for the regulation of TGF-β ligands, with specific components of the extracellular matrix known to segregate secreted cytokines to either inhibit TGF-β signalling or concentrate the ligands for future use (Horiguchi et al., 2012). Sequestration of TGF-β ligands was also demonstrated in *Drosophila* (Haerry, 2010) and proposed to facilitate autocrine TGF-β signalling in the larval NMJ (Kim and O’Connor, 2014). Further studies are required to firmly establish this mode of ligand control in regulating the *Drosophila* neuromuscular system. Considering the important role of TGF-β ligands in the regulation of muscle mass and NMJ function in mammalian model species (Fong et al., 2010; McLennan and Koishi, 1994) and in humans (Chen et al., 2016; McLennan and Koishi, 2002), these findings can point to novel mechanism for therapeutic interventions in pathologies associated with ageing and neuromuscular disorders.

## MATERIALS AND METHODS

### Fly stocks and husbandry

All stocks were maintained and all experiments were conducted at 25°C on a 12 h:12 h light:dark cycle at constant humidity using standard sugar/yeast/agar (SYA) media (15 g/l agar, 50 g/l sugar, 100 g/l autolyzed yeast, 100 g/l nipagin and 3 ml/l propionic acid) (Bass et al., 2007). Third instar wandering larvae used in the experiments were selected based on morphological (larval spiracles and mouth-hook) and behavioural criteria (location outside of the food). Tissue-specific expression was achieved with the GAL4-UAS system [GAL4-dependant upstream activator sequence] (Brand and Perrimon, 1993). Drosophila stocks used were: *UAS-miRNAmyo* (Awasaki et al., 2011), a gift from T. Awasaki from Tzumin Lee lab at Janelia Farm; *Mef2-GAL4* (Bloomington Stock Center #27390); UAS-PlumRNAi (Vienna *Drosophila* Resource Centre #101135) a gift from Oren Schuldiner lab at Weitzman Institute of Technology. The *Mef2-GAL4/UAS-miRNAmyo/ UAS-PlumRNAi* line was created using standard *Drosophila* crossing schemes. *w^Dah^* was the “wild-type” strain used in all experiments. The white Dahomey (*w^Dah^*) stock was derived by incorporation of the *w^1118^* mutation into the outbred Dahomey background by back-crossing.

### NMJ electrophysiology

Recordings were performed as described previously (Robinson et al., 2014). Two-electrode voltage clamp (TEVC) recordings using sharp electrodes were made from ventral longitudinal muscle 6 in abdominal segments 2-4 of wandering third instar larvae. NMJ recordings were performed using pClamp 10, an Axoclamp 900A amplifier and Digidata 1440A (Molecular Devices, USA) in haemolymph-like 3 (HL-3) solution: 70 mM NaCl, 5 mM KCl, 20 mM MgCl2, 10 mM NaHCO3, 115 mM sucrose, 5 mM trehalose, 5 mM HEPES and 2 mM CaCl2. Recording electrodes (10–30 MΩ) were filled with 3 M KCl. Miniature excitatory junctional currents (mEJCs) were recorded in the presence of 0.5 μM tetrodotoxin (Tocris, UK). All synaptic responses were recorded from muscles with input resistances ≥4 MΩ and resting potentials more negative than −60 mV at 25°C as differences in recording temperature cause changes in glutamate receptor kinetics and amplitudes (Postlethwaite et al., 2007). Holding potentials were −60 mV. Mean single eEJC amplitudes (stimulus: 0.1 ms, 1-5 V) are based on the mean peak eEJC amplitude in response to ten presynaptic stimuli (recorded at 0.2 Hz). Nerve stimulation was performed with an isolated stimulator (DS2A, Digitimer). The data were digitized at 10 kHz and for miniature recordings, 200 s recordings were analysed to obtain mean mEJC amplitudes and frequency values. mEJC and eEJC recordings were off-line low-pass filtered at 500 Hz and 1 kHz, respectively. Materials were purchased from Sigma-Aldrich (UK) unless otherwise stated.

### Immunocytochemistry and confocal microscopy

Immunocytochemistry and confocal microscopy were performed as described previously (Augustin et al., 2017) using Zeiss 700 inverted confocal microscope. All neuromuscular junction (NMJ) images and analyses were from NMJs on larval ventral longitudinal muscles 6 and 7 (hemi-segments A3-A4). For glutamate receptor (GluRIIA) and Brp staining, 3^rd^ instar larval preparations were dissected in modified HL-3 solution and fixed for 30 min in Bouin’s fixative. Mouse monoclonal anti-GluRIIA (8B4D2) and anti-Brp (nc82) antibodies were obtained from the University of Iowa Developmental Studies Hybridoma Bank (Iowa City, USA) and used at 1:100 and 1:20, respectively. AlexaFluor-conjugated goat anti-mouse secondary antibodies were used at 1:200. TRITC-labeled anti-horseradish peroxidase (HRP) antibody (staining neuronal membranes) was used at 1:100. To visualize larval muscles, phalloidin was added to fresh larval preparations fixed for 30 min with 4% paraformaldehyde. Measurements of the postsynaptic glutamate receptors were made using ImageJ by drawing a circle around individual GluRIIA clusters in type Ib synaptic boutons.

### Statistical analyses

Statistical analyses were performed using GraphPad Prism 5 software. For comparisons between two or more groups, a one-way ANOVA followed by a Tukey-Kramer test was used. In all instances, P<0.05 is considered statistically significant (*P<0.05; **P<0.01; ***P<0.001). Values are reported as the mean ± SEM. A 2-way ANOVA test was used to perform interaction calculations. The Kolmogorov-Smirnov (KS) test was used to analyse the cumulative distribution of mEJC amplitudes.

## ACKNOWLEDGEMENTS

We would like to thank Oren Schuldiner (Weizmann Institute of Science, Israel) for Plum lines and the Bloomington Drosophila Stock Center for reagents.

## REFERENCES

Augustin, H., McGourty, K., Steinert, J.R., Cocheme, H.M., Adcott, J., Cabecinha, M., Vincent, A., Halff, E.F., Kittler, J.T., Boucrot, E., et al. (2017). Myostatin-like proteins regulate synaptic function and neuronal morphology. Development 144, 2445–2455.

Augustin, H., and Partridge, L. (2009). Invertebrate models of age-related muscle degeneration. Biochimica et biophysica acta 1790, 1084–1094.

Awasaki, T., Huang, Y., O’Connor, M.B., and Lee, T. (2011). Glia instruct developmental neuronal remodeling through TGF-beta signaling. Nature neuroscience 14, 821–823.

Barker, D., and Ip, M.C. (1966). Sprouting and degeneration of mammalian motor axons in normal and de-afferentated skeletal muscle. Proc R Soc Lond B Biol Sci 163, 538–554.

Bass, T.M., Grandison, R.C., Wong, R., Martinez, P., Partridge, L., and Piper, M.D. (2007). Optimization of dietary restriction protocols in Drosophila. J Gerontol A Biol Sci Med Sci 62, 1071–1081.

Bezakova, G., and Ruegg, M.A. (2003). New insights into the roles of agrin. Nat Rev Mol Cell Biol 4, 295–308.

Boido, M., and Vercelli, A. (2016). Neuromuscular Junctions as Key Contributors and Therapeutic Targets in Spinal Muscular Atrophy. Front Neuroanat 10, 6.

Bombelli, F., Lispi, L., Porrini, S.C., Giacanelli, M., Terracciano, C., Massa, R., and Petrucci, A. (2016). Neuromuscular transmission abnormalities in myotonic dystrophy type 1: A neurophysiological study. Clin Neurol Neurosurg 150, 84–88.

Brand, A.H., and Perrimon, N. (1993). Targeted gene expression as a means of altering cell fates and generating dominant phenotypes. Development 118, 401–415.

Budnik, V. (1996). Synapse maturation and structural plasticity at Drosophila neuromuscular junctions. Current opinion in neurobiology 6, 858–867.

Butikofer, L., Zurlinden, A., Bolliger, M.F., Kunz, B., and Sonderegger, P. (2011). Destabilization of the neuromuscular junction by proteolytic cleavage of agrin results in precocious sarcopenia. FASEB J 25, 4378–4393.

Cappello, V., and Francolini, M. (2017). Neuromuscular Junction Dismantling in Amyotrophic Lateral Sclerosis. Int J Mol Sci 18.

Carnac, G., Vernus, B., and Bonnieu, A. (2007). Myostatin in the pathophysiology of skeletal muscle. Current genomics 8, 415–422.

Chand, K.K., Lee, K.M., Lee, J.D., Qiu, H., Willis, E.F., Lavidis, N.A., Hilliard, M.A., and Noakes, P.G. (2018). Defects in synaptic transmission at the neuromuscular junction precede motor deficits in a TDP-43(Q331K) transgenic mouse model of amyotrophic lateral sclerosis. FASEB J 32, 2676–2689.

Chen, J., Mizushige, T., and Nishimune, H. (2012). Active zone density is conserved during synaptic growth but impaired in aged mice. J Comp Neurol 520, 434–452.

Chen, J.L., Colgan, T.D., Walton, K.L., Gregorevic, P., and Harrison, C.A. (2016). The TGF-beta Signalling Network in Muscle Development, Adaptation and Disease. Adv Exp Med Biol 900, 97–131.

Clark, B.C., Mahato, N.K., Nakazawa, M., Law, T.D., and Thomas, J.S. (2014). The power of the mind: the cortex as a critical determinant of muscle strength/weakness. J Neurophysiol 112, 3219–3226.

Cruz-Jentoft, A.J., Baeyens, J.P., Bauer, J.M., Boirie, Y., Cederholm, T., Landi, F., Martin, F.C., Michel, J.P., Rolland, Y., Schneider, S.M., et al. (2010). Sarcopenia: European consensus on definition and diagnosis: Report of the European Working Group on Sarcopenia in Older People. Age Ageing 39, 412–423.

DeChiara, T.M., Bowen, D.C., Valenzuela, D.M., Simmons, M.V., Poueymirou, W.T., Thomas, S., Kinetz, E., Compton, D.L., Rojas, E., Park, J.S., et al. (1996). The receptor tyrosine kinase MuSK is required for neuromuscular junction formation in vivo. Cell 85, 501–512.

Delbono, O. (2000). Regulation of excitation contraction coupling by insulin-like growth factor-1 in aging skeletal muscle. J Nutr Health Aging 4, 162–164.

Delbono, O., O’Rourke, K.S., and Ettinger, W.H. (1995). Excitation-calcium release uncoupling in aged single human skeletal muscle fibers. J Membr Biol 148, 211–222.

Demontis, F., Piccirillo, R., Goldberg, A.L., and Perrimon, N. (2013). Mechanisms of skeletal muscle aging: insights from Drosophila and mammalian models. Dis Model Mech 6, 1339–1352.

DiAntonio, A., Petersen, S.A., Heckmann, M., and Goodman, C.S. (1999). Glutamate receptor expression regulates quantal size and quantal content at the Drosophila neuromuscular junction. The Journal of neuroscience : the official journal of the Society for Neuroscience 19, 3023–3032.

Drey, M., Sieber, C.C., Bauer, J.M., Uter, W., Dahinden, P., Fariello, R.G., Vrijbloed, J.W., and Fi, A.T.i.g. (2013). C-terminal Agrin Fragment as a potential marker for sarcopenia caused by degeneration of the neuromuscular junction. Exp Gerontol 48, 76–80.

Eddinger, T.J., Moss, R.L., and Cassens, R.G. (1985). Fiber number and type composition in extensor digitorum longus, soleus, and diaphragm muscles with aging in Fisher 344 rats. J Histochem Cytochem 33, 1033–1041.

Ezaki, T., Oki, S., Matsuda, Y., and Desaki, J. (2000). Age changes of neuromuscular junctions in the extensor digitorum longus muscle of spontaneous thymoma BUF/Mna rats. A scanning and transmission electron microscopic study. Virchows Arch 437, 388–395.

Fambrough, D.M., Drachman, D.B., and Satyamurti, S. (1973). Neuromuscular junction in myasthenia gravis: decreased acetylcholine receptors. Science 182, 293–295.

Featherstone, D.E., Rushton, E., and Broadie, K. (2002). Developmental regulation of glutamate receptor field size by nonvesicular glutamate release. Nature neuroscience 5, 141–146.

Featherstone, D.E., Rushton, E., Rohrbough, J., Liebl, F., Karr, J., Sheng, Q., Rodesch, C.K., and Broadie, K. (2005). An essential Drosophila glutamate receptor subunit that functions in both central neuropil and neuromuscular junction. The Journal of neuroscience : the official journal of the Society for Neuroscience 25, 3199–3208.

Ferns, M.J., Campanelli, J.T., Hoch, W., Scheller, R.H., and Hall, Z. (1993). The ability of agrin to cluster AChRs depends on alternative splicing and on cell surface proteoglycans. Neuron 11, 491–502.

Fong, S.W., McLennan, I.S., McIntyre, A., Reid, J., Shennan, K.I., and Bewick, G.S. (2010). TGF-beta2 alters the characteristics of the neuromuscular junction by regulating presynaptic quantal size. Proc Natl Acad Sci U S A 107, 13515–13519.

Gonzalez-Freire, M., de Cabo, R., Studenski, S.A., and Ferrucci, L. (2014). The Neuromuscular Junction: Aging at the Crossroad between Nerves and Muscle. Front Aging Neurosci 6, 208.

Gutmann, E., and Hanzlikova, V. (1973). Basic mechanisms of aging in the neuromuscular system. Mech Ageing Dev 1, 327–349.

Haerry, T.E. (2010). The interaction between two TGF-beta type I receptors plays important roles in ligand binding, SMAD activation, and gradient formation. Mechanisms of development 127, 358–370.

Herndon, L.A., Schmeissner, P.J., Dudaronek, J.M., Brown, P.A., Listner, K.M., Sakano, Y., Paupard, M.C., Hall, D.H., and Driscoll, M. (2002). Stochastic and genetic factors influence tissue-specific decline in ageing C. elegans. Nature 419, 808–814.

Hettwer, S., Dahinden, P., Kucsera, S., Farina, C., Ahmed, S., Fariello, R., Drey, M., Sieber, C.C., and Vrijbloed, J.W. (2013). Elevated levels of a C-terminal agrin fragment identifies a new subset of sarcopenia patients. Exp Gerontol 48, 69–75.

Horiguchi, M., Ota, M., and Rifkin, D.B. (2012). Matrix control of transforming growth factor-beta function. J Biochem 152, 321–329.

Kelly, S.S., and Robbins, N. (1983). Progression of age changes in synaptic transmission at mouse neuromuscular junctions. The Journal of physiology 343, 375–383.

Kim, M.J., and O’Connor, M.B. (2014). Anterograde Activin signaling regulates postsynaptic membrane potential and GluRIIA/B abundance at the Drosophila neuromuscular junction. PloS one 9, e107443.

Kong, L., Wang, X., Choe, D.W., Polley, M., Burnett, B.G., Bosch-Marce, M., Griffin, J.W., Rich, M.M., and Sumner, C.J. (2009). Impaired synaptic vesicle release and immaturity of neuromuscular junctions in spinal muscular atrophy mice. The Journal of neuroscience : the official journal of the Society for Neuroscience 29, 842–851.

Kreipke, R.E., Kwon, Y.V., Shcherbata, H.R., and Ruohola-Baker, H. (2017). Drosophila melanogaster as a Model of Muscle Degeneration Disorders. Curr Top Dev Biol 121, 83–109.

Lee, K.M., Chand, K.K., Hammond, L.A., Lavidis, N.A., and Noakes, P.G. (2017). Functional decline at the aging neuromuscular junction is associated with altered laminin-alpha4 expression. Aging (Albany NY) 9, 880–899.

Li, Y., Lee, Y., and Thompson, W.J. (2011). Changes in aging mouse neuromuscular junctions are explained by degeneration and regeneration of muscle fiber segments at the synapse. The Journal of neuroscience : the official journal of the Society for Neuroscience 31, 14910–14919.

Liebl, F.L., and Featherstone, D.E. (2008). Identification and investigation of Drosophila postsynaptic density homologs. Bioinform Biol Insights 2, 369–381.

Lo, P.C., and Frasch, M. (1999). Sequence and expression of myoglianin, a novel Drosophila gene of the TGF-beta superfamily. Mechanisms of development 86, 171–175.

Malatesta, M. (2012). Skeletal muscle features in myotonic dystrophy and sarcopenia: do similar nuclear mechanisms lead to skeletal muscle wasting? Eur J Histochem 56, e36.

Marquardt, J.U., Quasdorff, M., Varnholt, H., Curth, H.M., Mesghenna, S., Protzer, U., Goeser, T., and Nierhoff, D. (2011). Neighbor of Punc E11, a novel oncofetal marker for hepatocellular carcinoma. Int J Cancer 128, 2353–2363.

Mateos-Aierdi, A.J., Goicoechea, M., Aiastui, A., Fernandez-Torron, R., Garcia-Puga, M., Matheu, A., and Lopez de Munain, A. (2015). Muscle wasting in myotonic dystrophies: a model of premature aging. Front Aging Neurosci 7, 125.

McLennan, I.S., and Koishi, K. (1994). Transforming growth factor-beta-2 (TGF-beta 2) is associated with mature rat neuromuscular junctions. Neurosci Lett 177, 151–154.

McLennan, I.S., and Koishi, K. (2002). The transforming growth factor-betas: multifaceted regulators of the development and maintenance of skeletal muscles, motoneurons and Schwann cells. Int J Dev Biol 46, 559–567.

McPherron, A.C., Lawler, A.M., and Lee, S.J. (1997). Regulation of skeletal muscle mass in mice by a new TGF-beta superfamily member. Nature 387, 83–90.

Metter, E.J., Conwit, R., Metter, B., Pacheco, T., and Tobin, J. (1998). The relationship of peripheral motor nerve conduction velocity to age-associated loss of grip strength. Aging (Milano) 10, 471–478.

Mori, S., Koshi, K., and Shigemoto, K. (2014). The important role of the neuromuscular junction in maintaining muscle mass and strength. J Phys Fitness Sports Med 3, 111–114.

Nishimune, H., Badawi, Y., Mori, S., and Shigemoto, K. (2016). Dual-color STED microscopy reveals a sandwich structure of Bassoon and Piccolo in active zones of adult and aged mice. Sci Rep 6, 27935.

Oda, K. (1984). Age changes of motor innervation and acetylcholine receptor distribution on human skeletal muscle fibres. J Neurol Sci 66, 327–338.

Oliviero, A., Profice, P., Tonali, P.A., Pilato, F., Saturno, E., Dileone, M., Ranieri, F., and Di Lazzaro, V. (2006). Effects of aging on motor cortex excitability. Neurosci Res 55, 74–77.

Pansarasa, O., Rossi, D., Berardinelli, A., and Cereda, C. (2014). Amyotrophic lateral sclerosis and skeletal muscle: an update. Mol Neurobiol 49, 984–990.

Peron, S., Zordan, M.A., Magnabosco, A., Reggiani, C., and Megighian, A. (2009). From action potential to contraction: neural control and excitation-contraction coupling in larval muscles of Drosophila. Comparative biochemistry and physiology Part A, Molecular & integrative physiology 154, 173–183.

Petersen, S.A., Fetter, R.D., Noordermeer, J.N., Goodman, C.S., and DiAntonio, A. (1997). Genetic analysis of glutamate receptors in Drosophila reveals a retrograde signal regulating presynaptic transmitter release. Neuron 19, 1237–1248.

Piasecki, M., Ireland, A., Jones, D.A., and McPhee, J.S. (2016). Age-dependent motor unit remodelling in human limb muscles. Biogerontology 17, 485–496.

Plantie, E., Migocka-Patrzalek, M., Daczewska, M., and Jagla, K. (2015). Model organisms in the fight against muscular dystrophy: lessons from drosophila and Zebrafish. Molecules 20, 6237–6253.

Postlethwaite, M., Hennig, M.H., Steinert, J.R., Graham, B.P., and Forsythe, I.D. (2007). Acceleration of AMPA receptor kinetics underlies temperature-dependent changes in synaptic strength at the rat calyx of Held. The Journal of physiology 579, 69–84.

Pratt, S.J.P., Valencia, A.P., Le, G.K., Shah, S.B., and Lovering, R.M. (2015). Pre- and postsynaptic changes in the neuromuscular junction in dystrophic mice. Front Physiol 6, 252.

Proctor, D.N., Balagopal, P., and Nair, K.S. (1998). Age-related sarcopenia in humans is associated with reduced synthetic rates of specific muscle proteins. J Nutr 128, 351S–355S.

Qin, G., Schwarz, T., Kittel, R.J., Schmid, A., Rasse, T.M., Kappei, D., Ponimaskin, E., Heckmann, M., and Sigrist, S.J. (2005). Four different subunits are essential for expressing the synaptic glutamate receptor at neuromuscular junctions of Drosophila. The Journal of neuroscience : the official journal of the Society for Neuroscience 25, 3209–3218.

Querol, L., and Illa, I. (2013). Myasthenia gravis and the neuromuscular junction. Curr Opin Neurol 26, 459–465.

Rasse, T.M., Fouquet, W., Schmid, A., Kittel, R.J., Mertel, S., Sigrist, C.B., Schmidt, M., Guzman, A., Merino, C., Qin, G., et al. (2005). Glutamate receptor dynamics organizing synapse formation in vivo. Nature neuroscience 8, 898–905.

Robinson, S.W., Nugent, M.L., Dinsdale, D., and Steinert, J.R. (2014). Prion protein facilitates synaptic vesicle release by enhancing release probability. Human molecular genetics.

Rudolf, R., Khan, M.M., Labeit, S., and Deschenes, M.R. (2014). Degeneration of neuromuscular junction in age and dystrophy. Front Aging Neurosci 6, 99.

Ruiz-Canada, C., and Budnik, V. (2006). Introduction on the use of the Drosophila embryonic/larval neuromuscular junction as a model system to study synapse development and function, and a brief summary of pathfinding and target recognition. Int Rev Neurobiol 75, 1–31.

Salbaum, J.M., and Kappen, C. (2000). Cloning and expression of nope, a new mouse gene of the immunoglobulin superfamily related to guidance receptors. Genomics 64, 15–23.

Shigemoto, K., Kubo, S., Mori, S., Yamada, S., Akiyoshi, T., and Miyazaki, T. (2010). Muscle weakness and neuromuscular junctions in aging and disease. Geriatr Gerontol Int 10 Suppl 1, S137–147.

Shin, J., Tajrishi, M.M., Ogura, Y., and Kumar, A. (2013). Wasting mechanisms in muscular dystrophy. Int J Biochem Cell Biol 45, 2266–2279.

Sigrist, S.J., Reiff, D.F., Thiel, P.R., Steinert, J.R., and Schuster, C.M. (2003). Experience-dependent strengthening of Drosophila neuromuscular junctions. The Journal of neuroscience : the official journal of the Society for Neuroscience 23, 6546–6556.

Sigrist, S.J., Thiel, P.R., Reiff, D.F., and Schuster, C.M. (2002). The postsynaptic glutamate receptor subunit DGluR-IIA mediates long-term plasticity in Drosophila. The Journal of neuroscience : the official journal of the Society for Neuroscience 22, 7362–7372.

Takahashi, K.F., Kiyoshima, T., Kobayashi, I., Xie, M., Yamaza, H., Fujiwara, H., Ookuma, Y., Nagata, K., Wada, H., Sakai, T., et al. (2010). Protogenin, a new member of the immunoglobulin superfamily, is implicated in the development of the mouse lower first molar. BMC Dev Biol 10, 115.

Tintignac, L.A., Brenner, H.R., and Ruegg, M.A. (2015). Mechanisms Regulating Neuromuscular Junction Development and Function and Causes of Muscle Wasting. Physiol Rev 95, 809–852.

Wagh, D.A., Rasse, T.M., Asan, E., Hofbauer, A., Schwenkert, I., Durrbeck, H., Buchner, S., Dabauvalle, M.C., Schmidt, M., Qin, G., et al. (2006). Bruchpilot, a protein with homology to ELKS/CAST, is required for structural integrity and function of synaptic active zones in Drosophila. Neuron 49, 833–844.

Wigg, K.G., Feng, Y., Crosbie, J., Tannock, R., Kennedy, J.L., Ickowicz, A., Malone, M., Schachar, R., and Barr, C.L. (2008). Association of ADHD and the Protogenin gene in the chromosome 15q21.3 reading disabilities linkage region. Genes Brain Behav 7, 877–886.

Wokke, J.H., Jennekens, F.G., van den Oord, C.J., Veldman, H., Smit, L.M., and Leppink, G.J. (1990). Morphological changes in the human end plate with age. J Neurol Sci 95, 291–310.

Wong, Y.H., Lu, A.C., Wang, Y.C., Cheng, H.C., Chang, C., Chen, P.H., Yu, J.Y., and Fann, M.J. (2010). Protogenin defines a transition stage during embryonic neurogenesis and prevents precocious neuronal differentiation. The Journal of neuroscience : the official journal of the Society for Neuroscience 30, 4428–4439.

Xiang, Y., Yang, T., Pang, B.Y., Zhu, Y., and Liu, Y.N. (2016). The Progress and Prospects of Putative Biomarkers for Liver Cancer Stem Cells in Hepatocellular Carcinoma. Stem Cells Int 2016, 7614971.

Xu, R., and Salpeter, M.M. (1997). Acetylcholine receptors in innervated muscles of dystrophic mdx mice degrade as after denervation. The Journal of neuroscience : the official journal of the Society for Neuroscience 17, 8194–8200.

Young, A., Stokes, M., and Crowe, M. (1985). The size and strength of the quadriceps muscles of old and young men. Clin Physiol 5, 145–154.

Yu, X.M., Gutman, I., Mosca, T.J., Iram, T., Ozkan, E., Garcia, K.C., Luo, L., and Schuldiner, O. (2013). Plum, an immunoglobulin superfamily protein, regulates axon pruning by facilitating TGF-beta signaling. Neuron 78, 456–468.

